# Reducing effects of particle adsorption to the air-water interface in cryoEM

**DOI:** 10.1101/288340

**Authors:** Alex J. Noble, Hui Wei, Venkata P. Dandey, Zhening Zhang, Yong Zi Tan, Clinton S. Potter, Bridget Carragher

**Author notes:** These authors contributed equally to this work (Alex J. Noble, Hui Wei, Venkata P. Dandey).

## Abstract

Most protein particles prepared in vitreous ice for single particle cryo-electron microscopy are adsorbed to air-water or substrate-water interfaces, potentially causing particles to adopt preferred orientations. Using the Spotiton robot and nanowire grids, we can reduce some of the deleterious effects of the air-water interface by decreasing the dwell time of particles in thin liquid films. We demonstrate this by using single particle cryoEM and cryoET on three biological samples.

---

Single particle cryo-electron microscopy (cryoEM) allows for the structural study of purified proteins in solution to near-atomic resolutions^1,2^. Proteins are preserved in their hydrated state by spreading the sample out in a thin layer of buffer solution supported on a cryoEM grid and rapidly plunging the grid into a cryogen so as to convert the liquid layer into vitreous ice^3,4^. Alignment, classification, and reconstruction of a sufficient number of EM images of randomly oriented protein particles provides 3D density maps. Advances in electron microscope hardware, cameras, and image processing methods have enabled cryoEM as a method for reconstructing a wide range of protein complexes to near-atomic resolution in near-native conditions in multiple functional states^5,6^.

Tomographic studies of a wide range of single particle samples have shown that the vast majority of proteins prepared over holey substrates using standard vitrification methods are adsorbed to air-water interfaces^7^. This has the potential to cause the protein particles to adopt preferred orientations as well as the possibility of damaging or degrading the protein structure^7,8^. There are various options for avoiding contact with the air-water interface and its potential deleterious effects, including using surfactants as a barrier^8^, sequestering the particles to a support film^8^, or by outrunning some of the surface effects by reducing the length of time that the sample dwells in the thin liquid film prior to vitrification. It is this latter approach that we describe here; if the dwell time is reduced sufficiently, the particles in solution may either not have time to diffuse completely to the air-water or substrate-water interfaces or may not have fully equilibrated after having arrived there, depending on their affinity for the interfaces.

For most cryoEM vitrification devices (FEI Vitrobot, Gatan CP3, Leica EM GP, manual plungers) the time that elapses between the wicking of a sample to a thin film and the grid entering the cryogen is typically on the order of 1 second or greater. Assuming a thin film of thickness ~100nm, various estimates indicate that the protein particles will collide with an air-water interface about 100-1000 times during this time interval^9,10^, providing ample opportunity for adsorption and preferential orientation. There are at least three devices currently under development that allow for much more rapid plunge-freezing: a microfluidic spray-plunging machine developed by the Frank group^11^, a surface acoustic wave based microfluidic dispenser by the de Marco group^12^, and Spotiton, a robotic device that dispenses picoliter-volumes of sample onto a self-blotting nanowire grid as it flies past *en route* to vitrification^13–16^. Here we are using Spotiton to achieve rapid plunge times, however the methods are completely generalizable and we expect to see many new devices focus on providing faster plunge times as an option. We call the time interval between sample application to the grid and vitrification the spot-to-plunge time. Previously, the spot-to-plunge times on the Spotiton robot were on the order of 500 ms or more, but we recently modified the device to achieve spot-to-plunge times on the order of 100 ms.

Here we demonstrate, using three different specimens (hemagglutinin, insulin receptor bound to insulin^17^, and apoferritin), that by decreasing the spot-to-plunge time and thus reducing the dwell time of the sample in the thin liquid layer, the orientations of particles adsorbed to the air-water interfaces may be increased, and the density of non-adsorbed particles in grid holes may be increased. We used Spotiton to vitrify samples with spot-to-plunge times of 100-200 ms and compared these results, using cryoEM and cryoET^7^, to those with spot-to-plunge times of 400 ms to ~1 s, serving as controls.

With longer spot-to-plunge times (500 ms to ~1 s), hemagglutinin exhibits pronounced preferred orientation, presenting very few side-views of the particles in 2D class averages (Fig. 1a), while 2D class averages of insulin-bound insulin receptor provide a limited set of particle orientations (Fig. 1b). For both samples, the vast majority of particles are closely associated with the air-water interfaces (**Supplementary Videos 1 & 2**). As a consequence of the preferred orientation, coherent initial models of hemagglutinin and insulin receptor could not be generated and isotropic 3D reconstructions could not be obtained unless micrographs were acquired with the grid tilted relative to the electron beam^18^, which however imposes collection, processing, and resolution limitations. Apoferritin with 0.5 mM TCEP (15 mg/mL = 20,530 particles/μm^3^ in solution), when plunged with a longer spot-to-plunge time of 500 ms, primarily adsorbs to the air-water interfaces, although its high symmetry and the prevalence of local tilt in exposure areas^7^ effectively negate any issues of preferred orientation (Fig. 2a, **Supplementary Video 3**). The volume of ice occupied by the non-adsorbed apoferritin particles contains a particle density of about 1,668 particles/μm^3^ (Supplementary Fig. 1).

**Figure 1.**
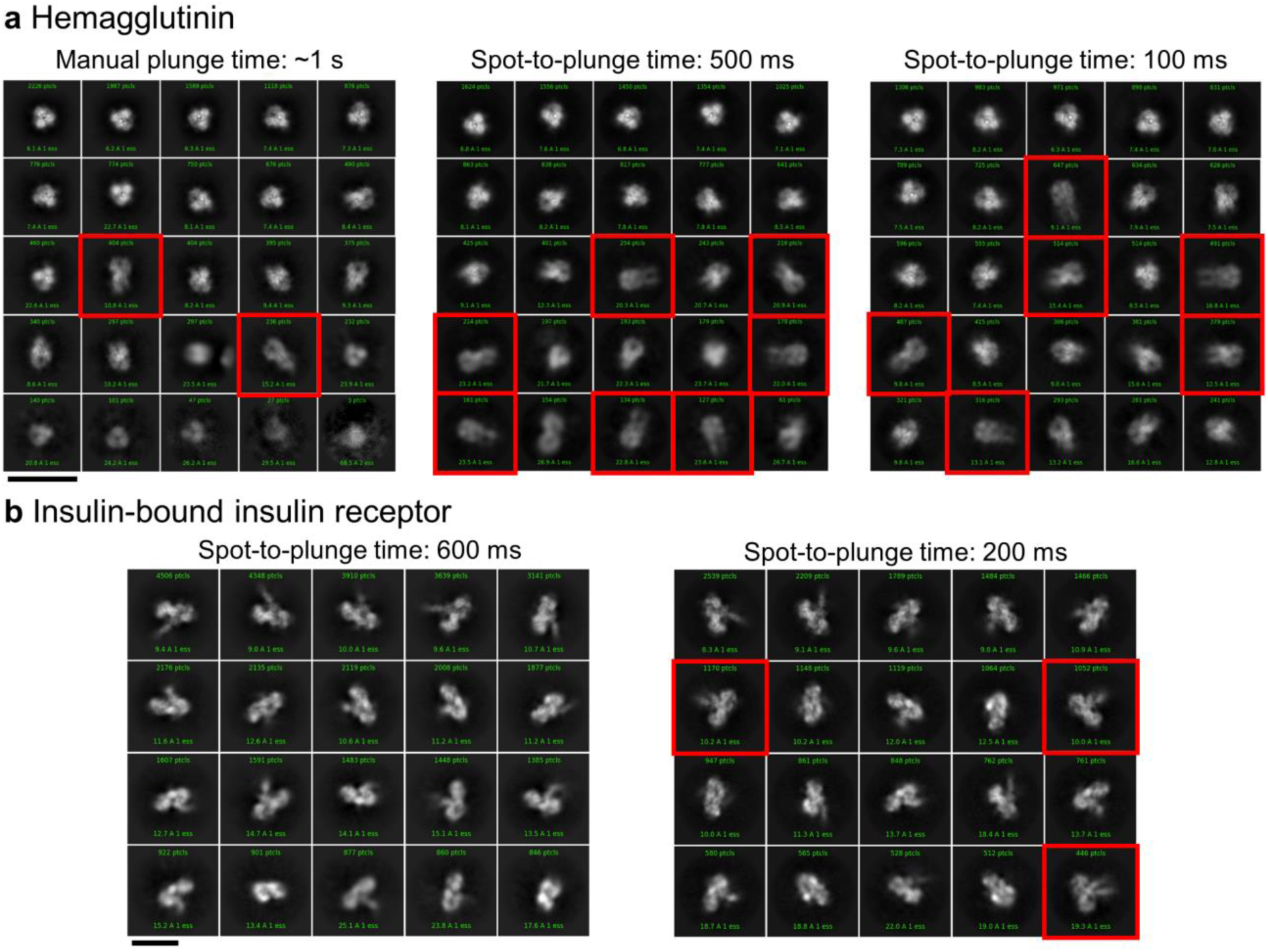
Single particle cryoEM 2D classification comparison between long and short spot-to-plunge times. (a) 2D classification of hemagglutinin plunged with a Gatan CP3 with an estimated blot-to-plunge time of ~1 s shows severe preferred orientation in the resulting 2D class averages, where only 4% of classes are side-views (red squares) (left). When plunged with a spot-to-plunge time of 500 ms, the percentage of side-views increases to 9% (middle). When plunged with a spot-to-plunge time of 100 ms, 19% of classes are side-views (right). (b) 2D classification of insulin-bound insulin receptor with a spot-to-plunge time of 600 ms (left) shows a well-populated, yet incomplete, set of particle views in the class averages. When plunged with a spot-to-plunge time of 200 ms, several additional side-views are recovered (red squares). Scale bars are 20 nm.

A shorter spot-to-plunge time of 100 ms for hemagglutinin results in much reduced preferred orientations (Fig. 1a), and results in a 3D reconstructed map at 3.8 Å (Supplementary Fig. 2) that is more isotropic and better resolved compared to the 4.2 Å map^18^ produced using grid tilt. Similarly, for the insulin-bound insulin receptor with a spot-to-plunge time of 200 ms, the reduced preferred orientation provides additional critical views of the complex (Fig. 1b), producing a 4.9 Å 3D reconstruction (Supplementary Fig. 3) that is of higher quality and more isotropic than that derived from images of tilted grids^17^. While the preferred orientation of hemagglutinin and insulin receptor were each significantly reduced with shorter spot-to-plunge times, the majority of particles still remained adsorbed to the air-water interfaces (**Supplementary Videos 4 & 5**). However, in the case of apoferritin with TCEP, a spot-to-plunge time of 170 ms vs. 500 ms significantly increased the density of non-adsorbed particles (Fig. 2b, **Supplementary Video 6**). The density of particles not adsorbed to the air-water interfaces increased by a factor of ~20x to about 31,725 particles/μm^3^ (Supplementary Fig. 4). When apoferritin at a lower concentration was prepared without TCEP (6 mg/mL = 8,212 particles/μm^3^ in solution), the density of non-adsorbed particles increased from about 3,043 particles/μm^3^ with a spot-to-plunge time of 500 ms to about 17,927 particles/μm^3^ with a spot-to-plunge time of 100 ms (Supplementary Figs. 5 & 6, **Supplementary Videos 7 & 8**). We note that there are far fewer particles adsorbed to the air-water interface for the apoferritin sample with TCEP added to the buffer but it is also likely that additional factors are contributing to the observed differences. These factors include effects of evaporation, estimated at 300 Å/s for (~85% relative humidity, ~70 °F). A full understanding of these and other effects clearly requires much further study under well-defined and controlled conditions.

**Figure 2.**
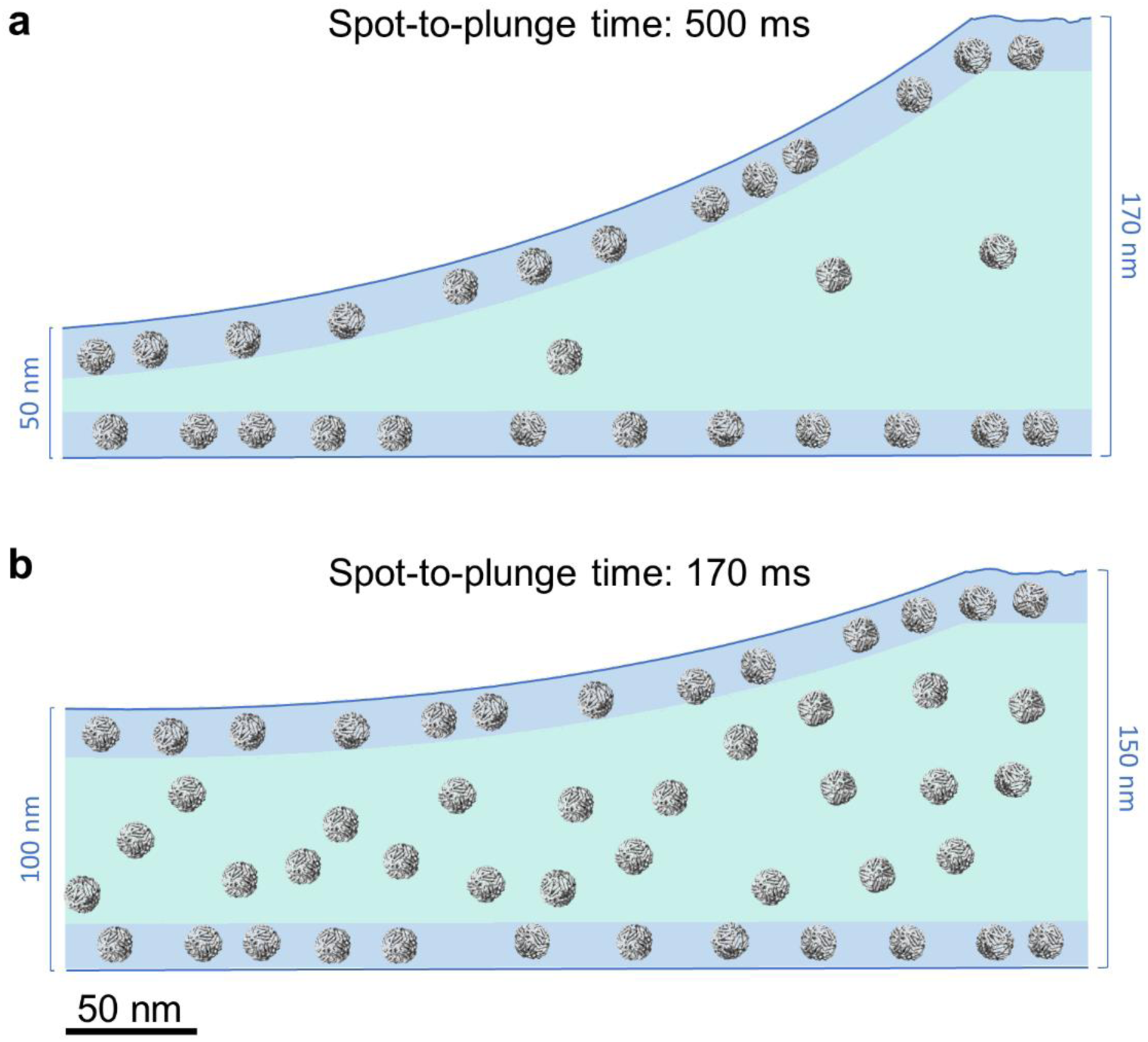
CryoET cross-sectional depictions of apoferritin comparing long and short spot-to-plunge times. Areas colored in blue represent locations where particles are adsorbed to the air-water interface while areas in teal represent the volume in between. (a) Apoferritin with 0.5 mM TCEP (15 mg/mL = 20,530 particle/μm^3^) plunged with a spot-to-plunge time of 500 ms shows that the vast majority of particles are adsorbed to air-water interfaces; the density of free-floating particles in the volume of ice is about 1,668 particles/μm^3^. (b) Apoferritin plunged with a spot-to-plunge time of 170 ms shows that while many particles are still adsorbed to air-water interfaces, the density of free-floating particles in the volume of ice increased by about 20x to 31,725 particles/μm^3^.

These three example specimens show that deleterious effects of particle adsorption to air-water interfaces can be considerably reduced by decreasing the time between sample application and plunge-freezing, thus increasing the density of non-adsorbed particles in holes and reducing the opportunities for particles to equilibrate at air-water interfaces. We anticipate that the interface effects can be further reduced by decreasing the spot-to-plunge time even further, to a few 10s of msecs. The observations in this study do not directly address whether speeding up the plunging results in fewer particles being adsorbed to the air-water interface, but do clearly indicate that the overall effect is to reduce preferred orientation. In the case of Spotiton, faster spot-to-plunge times may be more challenging for controlling ice thickness and may require higher mesh grids (to increase nanowire density) or more accurate control of relative humidity. However, while thinner ice is usually the ideal outcome, we note that near-atomic resolution structures are attainable in ice thicker than 100 nm^19^. Finally, if the density of non-adsorbed particles can be significantly increased, performing per-particle CTF estimation^20–22^ on these images provides the possibility for *in silico* identification of non-adsorbed vs. adsorbed particles in single particle cryoEM. As a result, it may be possible to derive single particle cryoEM structures based only on non-adsorbed particles, thus allowing for more explicit studies of the effects of absorption on structure and preferred orientation.

## METHODS

Methods, including statements of data availability and any associated accession codes and references, are available in the online version of the paper.

## ACKNOWLEDGEMENTS

The authors wish to thank Prof. Robert Glaeser at Lawrence Berkeley National Laboratory for helpful discussions, Dr. Kotaro Kelley at the New York Structural Biology Center (NYSBC) for apoferritin purification, and the Electron Microscopy Group at NYSBC for microscope calibration and assistance.

A.J.N. was supported by a grant from the NIH National Institute of General Medical Sciences (F32GM128303). Y.Z.T. was supported in part by the Agency for Science, Technology and Research Singapore. This work was performed at the Simons Electron Microscopy Center and National Resource for Automated Molecular Microscopy located at the New York Structural Biology Center, supported by grants from the Simons Foundation (SF349247), NYSTAR, and the NIH National Institute of General Medical Sciences (GM103310) with additional support from the Agouron Institute (F00316) and NIH (OD019994).

## AUTHOR CONTRIBUTIONS

A.J.N. collected and aligned tilt-series, and analyzed cryoET data. H.W. and V.P.D. performed the Spotiton experiments and the cryoEM experiments. H.W. made nanowire grids and collected tilt-series. H.W., V.P.D., and Y.Z.T. analyzed single particle cryoEM data. A.J.N., H.W., V.P.D., Z.Z., C.S.P., and B.C. conceived and designed the experiments. A.J.N. and B.C. wrote the manuscript.

## ONLINE METHODS

### Sample preparation

#### Hemagglutinin

Hemagglutinin was prepared as described in Tan, Y. Z. et al., 2017^18^.

#### Insulin-bound insulin receptor

Insulin-bound insulin receptor was prepared as described in Scapin, G. et al., 2018^17^.

#### Apoferritin

Apoferritin from equine spleen (Sigma-Aldrich) as shown in Figure 2, Supplementary Figures 1 & 4, and **Supplementary Videos 3 & 6** was prepared by diluting the stock sample to 15 mg/mL with 0.5 mM TCEP.

Apoferritin from equine spleen (Sigma-Aldrich) as shown in Supplementary Figures 5 & 6 and **Supplementary Videos 7 & 8** was prepared as follows. 100 μL of sample at 25 mg/mL was diluted with 1 mL QA, 1 mL was loaded into a Q column, eluted with QB, and the main peak was pooled and concentrated to 0.5 mL. This 0.5 mL was then loaded onto a 10300 S200 column (GE) in QF, eluted at approximately 10.5 mL, and concentrated to working condition. [QA = 20 mM HEPES at pH 7.5, 0.1 mM EDTA at pH 8.0, 1 mM DTT; QB = QA + 1.5 M NaCl; QF = 150 mM NaCl, 20 mM HEPES at pH 7.5]

#### Grid preparation and vitrification

Nanowire grids were prepared as described in Razinkov et al., 2016^14^ and Wei et al., 2018^16^.

The samples were vitrified using the semi-automated Spotiton V1.0 robot, a novel device for preparing cryoEM samples using piezo dispensing to apply small (50 pL) drops of sample across a "self-blotting” nanowire grid as it flies past *en route* to plunge into liquid ethane. Nanowire grids, manufactured in-house, backed by lacey carbon film supports were used for all experiments. Nanowire grids were plasma cleaned for 10 secs (O_2_ + H_2_) using a Solarus 950 (Gatan, Inc.). Sample was dispensed onto a grid dropping vertically past the dispense head in 50 pL drops for a total of ~5 nL of sample applied as a stripe across the grid which was then plunged into liquid ethane. The time between sample application to the grid and plunging into liquid ethane (spot-to-plunge time) ranged from 100 to 800 ms. Spotiton was operated at ~85% relative humidity and ambient temperature (~70 °F). Under these conditions, evaporation is estimated to be 300 Å/s.

The time of flight of the 50 pL drops from the nozzle to the grid is estimated to be 0.5 ms and only the first drop has time to form a "skin” as it waits in the nozzle. Thus, this phase is likely unimportant in the formation of the air-water interface. After the first contact of drops to grid, the drops have a fairly large volume relative to surface area and spread out under their own momentum. Contact with the nanowires results in very rapid formation of a thin layer, estimated to occur within ~20 ms. We thus believe that most of the air-water interface effects occur in the time of flight between the formation of the thin layer and the plunge into ethane.

### Single particle cryoEM data collection

#### Hemagglutinin

Single particle micrographs were collected on a Titan Krios (Thermo Fischer Scientific) equipped with an energy filter, Cs corrector, and a K2 counting camera (Gatan, Inc.); the microscope was operated at 300 kV at a nominal magnification of 130,000x, with a calibrated pixel size of 1.061 Å. Exposure was set to 10 secs (40 frames/image), for a total dose of 73.24 e^−^/Å^2^ with a defocus range of 1 to 2.1 μm. A total of 896 images were collected in three sessions using Leginon^23^.

#### Insulin-bound insulin receptor

Single particle micrographs were collected on a Titan Krios (Thermo Fischer Scientific) equipped with an energy filter and a K2 BioQuantum counting camera (Gatan, Inc.); the microscope was operated at 300 kV at a nominal magnification of 105,000x, with a calibrated pixel size of 1.096 Å. Exposure was set to 10 secs (40 frames/image), for a total dose of 66 e^−^ /Å^2^ with a defocus range of 1 to 2.5 μm. Collection was performed over three sessions using Leginon. For the 200 ms spot-to-plunge experiment, a total of 1,526 micrographs were used for single particle processing while 1,866 micrographs were used for the 800 ms spot-to-plunge experiment.

### Single particle cryoEM data processing

#### Hemagglutinin

Frames were aligned using MotionCor2^24^; global and per-particle CTF was estimated using gCTF^20^. Particle picking was performed using Gautomatch (http://www.mrc-lmb.cam.ac.uk/kzhang/) or DoG Picker^25^ followed by one round of 2D classification to remove false picks. A total of 104,365 particles were used for the final, symmetrized homogeneous 3D refinement in CryoSPARC^26^, producing a 3.77 Å map as shown in Supplementary Figure 2a.

#### Hemagglutinin with CP3

The hemagglutinin dataset EMPIAR-10096 plunged with a Gatan CP3 from the paper Tan, Y. Z. et al., 2017^18^ was used. From the 130,000 particle stack, a total of 15,000 particles were randomly selected for 2D classification and 3D refinement.

#### Insulin-bound insulin receptor

Micrograph frames were aligned using MotionCor2^24^; global and per-particle CTF was estimated using gCTF^22^. Particle picking was performed using Gautomatch (http://www.mrc-lmb.cam.ac.uk/kzhang/) followed by one round of 2D refinement to remove the false picks. Data processing from 2D classification through final reconstruction was performed using CryoSPARC as described in Scapin et al., 2018^17^. The completeness of the maps was assessed the Euler angle orientation distribution as calculated by CryoSPARC and by evaluating each map’s directional FSC.

For the 200 ms spot-to-plunge experiment, a total of 83,513 particles were selected for ab-initio 3D classification from which the best 40,390 particles were used for the final, symmetrized homogeneous 3D refinement, producing a 4.93 Å map.

For the 800 ms spot-to-plunge experiment, a total of 276,466 particles were selected for ab-initio 3D classification from which the best 70,276 particles were used for the final, symmetrized homogeneous 3D refinement, producing a 6.09 Å map.

Tilt-series data collection. Tilt-series were collected on a Titan Krios (Thermo Fischer Scientific) equipped with an energy filter and a K2 counting camera (Gatan, Inc.) and on a Tecnai F20 (Thermo Fischer Scientific) with a DE-20 camera (Direct Electron). Most tilt-series were collected nominally from −45° to 45° with 3° fixed or Saxton scheme^27^ increments using either Leginon^23,28^ or SerialEM^29^ with a nominal defocus near 5 microns, per-tilt image doses between 2 and 3 e-/Å^2^, and a pixel size of 2.16 Å (K2) and 2.34 Å (DE-20). K2 tilt images were whole-frame aligned using MotionCor2.

### Tilt-series data processing

Tilt-series were aligned with Appion-Protomo^30,31^ by first dose compensating the images with respect to the total accumulated dose^32^ of the tilt-series, coarse aligning tilt images, manually fixing coarse alignment if necessary, refining tilt-series alignment over several dozen iterations, and reconstructing with Tomo3D SIRT^33,34^. Tilt-series were not CTF corrected.

### Tomogram particle picking

Particles in the apoferritin tomograms (**Supplementary Videos 3, 6, 7, & 8**) were picked manually using Dynamo^35^. Particle picks were separated by location: 1) Particles over the carbon in the zero-degree projection direction or adsorbed to the carbon, 2) Particles adsorbed to the air-water interfaces in the grid holes, and 3) Non-adsorbed particles in the grid holes. The particle picks and the resulting particle densities are shown in Supplementary Figures 1, 4, 5, & 6. Partial particles (where at least half of the particle is visible) were picked. The density of adsorbed particles was calculated by estimating the 2D surface area of each air-water interface and multiplying by the diameter of apoferritin to obtain the volume. We estimate that particle identification is accurate to within 1% and is complete to within 1% while volume measurements are estimated to be accurate to within 5% due to ice curvature and grid shape approximations, thus density calculations are likely accurate to within 5%. Density calculations were only performed on one tomogram (one location on the grid) per apoferritin grid, therefore while we estimate our counting accuracy for this one tomogram to be 5% it is clearly not applicable across the entire grid.

### Data availability statement

Single particle half maps, full sharpened maps, and masks for insulin-bound insulin receptor (200 ms spot-to-plunge time), insulin-bound insulin receptor (800 ms), and hemagglutinin (100 ms) have been deposited to the Electron Microscopy Data Bank (EMDB) with accession codes EMD-7788, EMD-7791, and EMD-7792, respectively. The full single particle collection of hemagglutinin (100 ms and 500 ms) has been deposited to the Electron Microscopy Pilot Image Archive (EMPIAR) with accession codes EMPIAR-10175 and EMPIAR-10197, respectively. Single particle cryoET tomograms have been deposited to the EMDB with accession codes EMD-7623, EMD-7624, EMD-7625, EMD-7150, EMD-7627, EMD-7628, EMD-7629, and EMD-7630. Single particle cryoET tilt-series, cryoET tilt-series alignment runs with Appion-Protomo, cryoET tomograms, and apoferritin particle picks have been deposited to the EMPIAR with accession codes EMPIAR-10169, EMPIAR-10170, EMPIAR-10171, EMPIAR-10141, EMPIAR-10172, EMPIAR-10129, EMPIAR-10173, and EMPIAR-10174.

---

**Supplementary Figure 1.**
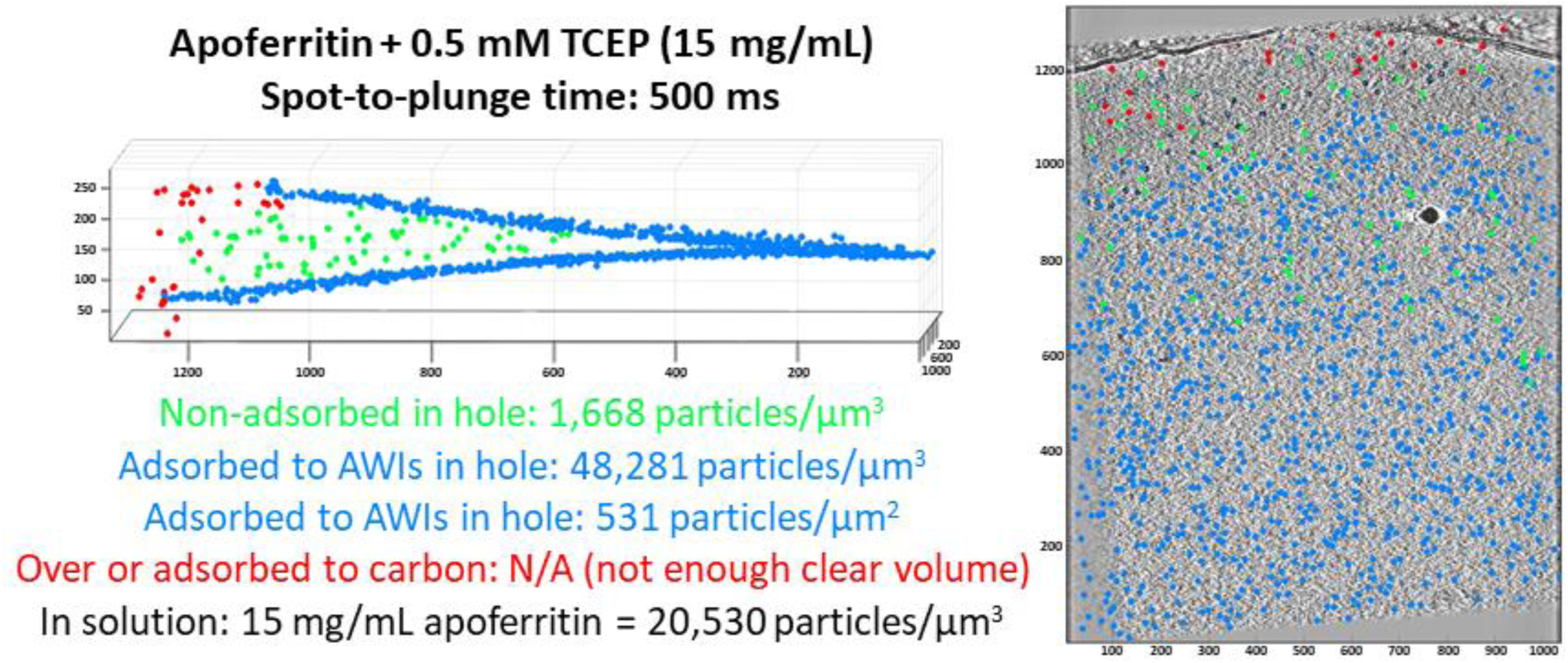
Particle distribution and concentrations of apoferritin (15 mg/mL = 20,530 particles/μm^3^ in solution) with 0.5 mM TCEP prepared using Spotiton and nanowire grids with spot-to-plunge time of 500 ms. Particle picking and depictions were performed in Dynamo^35^. Left depiction is a side-view aligned roughly perpendicular to the air-water interfaces. Right depiction is a top-down view with a tomographic slice included. Particles were manually picked according to their estimated locations relative to the air-water interface and carbon. Calculated particle densities (particles/μm^3^) are: 1,668 non-adsorbed in the hole and 48,281 adsorbed to the air-water interfaces. Particle surface density at the air-water interface is 531 particles/μm^2^. Particle concentration calculations are estimated to be accurate to within 5%. Scale: 100 pixels is ~94 nm.

**Supplementary Figure 2.**
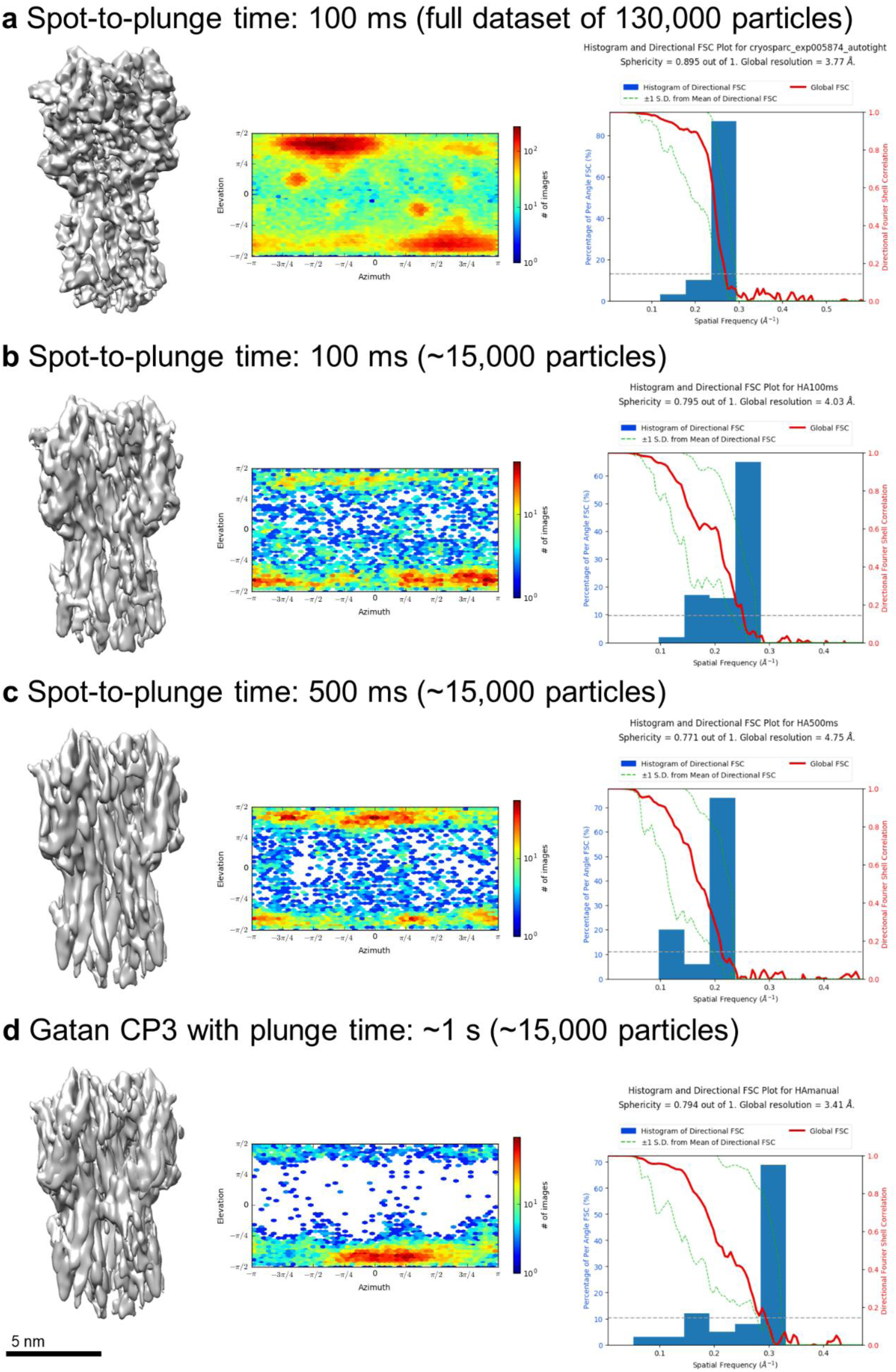
3D reconstructions of hemagglutinin with varying plunge times. From left to right: 3D reconstruction of hemagglutinin (C3 symmetry applied) from the given number of particles, orientation direction distribution plot (C1 symmetry applied), directional FSC plot (C3 symmetry applied). The directional FSC plots are created by calculating the FSC from 100 solid angles as a way to account for orientation when reporting resolution^18^; the histogram shows how many solid angles report a resolution in a particular range, which is reported as ‘Percentage of Per Angle FSC (%)’. (a) The full 100 ms spot-to-plunge time dataset of ~130,000 particles shows a relatively isotropic distribution, a sphericity of 0.895, and a resolution of 3.77 Å. (b) ~15,000 particles from the 100 ms spot-to-plunge time dataset (normalized to the number of particles available in the 500 ms dataset in (c)) shows a somewhat isotropic distribution, a sphericity of 0.795, and a resolution of 4.03 Å. (c) ~15,000 particles from the 500 ms spot-to-plunge time dataset shows a somewhat isotropic distribution, a sphericity of 0.771, and a resolution of 4.75 Å. (d) ~15,000 particles from the CP3-plunged sample shows a severely anisotropic distribution, an erroneous sphericity of 0.794, and an erroneous resolution of 3.41 Å.

**Supplementary Figure 3.**
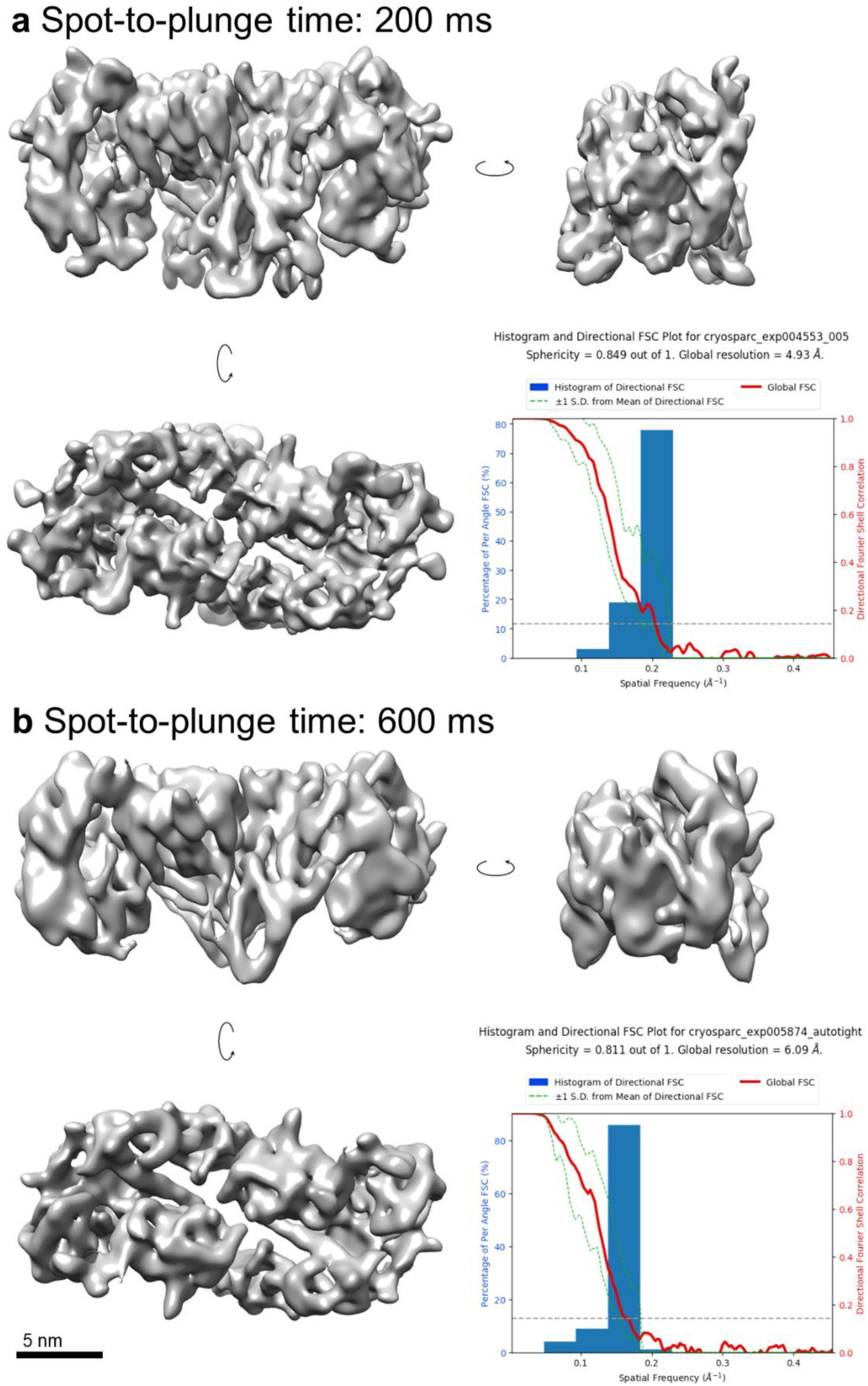
Single particle cryoEM of insulin-bound insulin receptor with spot-to-plunge times of 200 ms (a) and 600 ms (b). As shown in Figure 1, some side-views of insulin receptor are missing in the class averages of the 600 ms spot-to-plunge preparation. This resulted in a reduced sphericity of the directional FSC (0.811 vs. 0.849 for the 200 ms spot-to-plunge time) and worse resolution in the calculated map (6.09 Å vs. 4.93 Å for 200 ms). The greater anisotropy associated with for the longer spot-to-plunge time is reflected in the stretching artifacts in the 3D reconstruction.

**Supplementary Figure 4.**
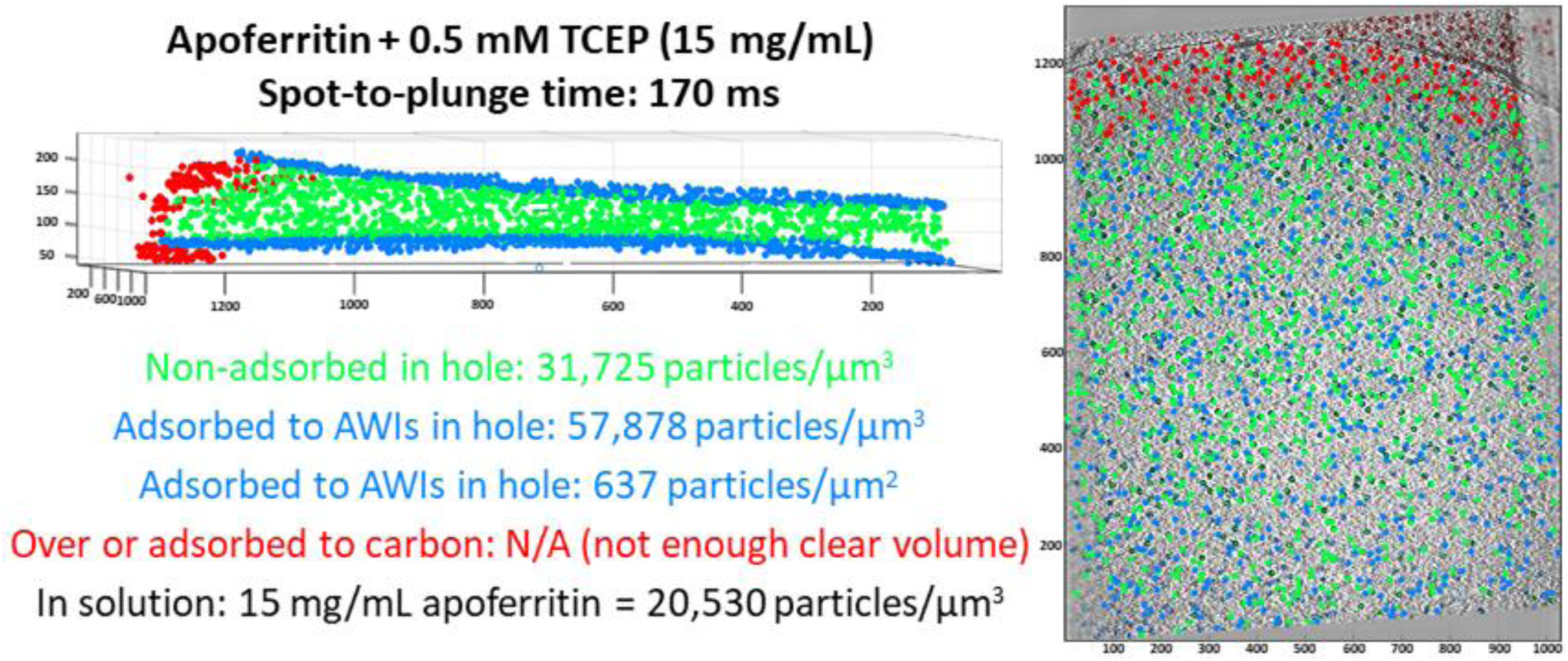
Particle distribution and concentrations of apoferritin (15 mg/mL = 20,530 particles/μm^3^ in solution) with 0.5 mM TCEP prepared using Spotiton and nanowire grids with spot-to-plunge time of 170 ms. Left depiction is a side-view aligned roughly perpendicular to the air-water interfaces. Right depiction is a top-down view with a tomographic slice included. Particles were manually picked according to their estimated locations relative to the air-water interface and carbon. Calculated particle densities (particles/μm^3^) are: 31,725 in the hole and 57,878 adsorbed to the air-water interfaces. Particle surface density at the air-water interface is 637 particles/μm^2^. Particle concentration calculations are estimated to be accurate to within 5%. Scale: 100 pixels is ~94 nm.

**Supplementary Figure 5.**
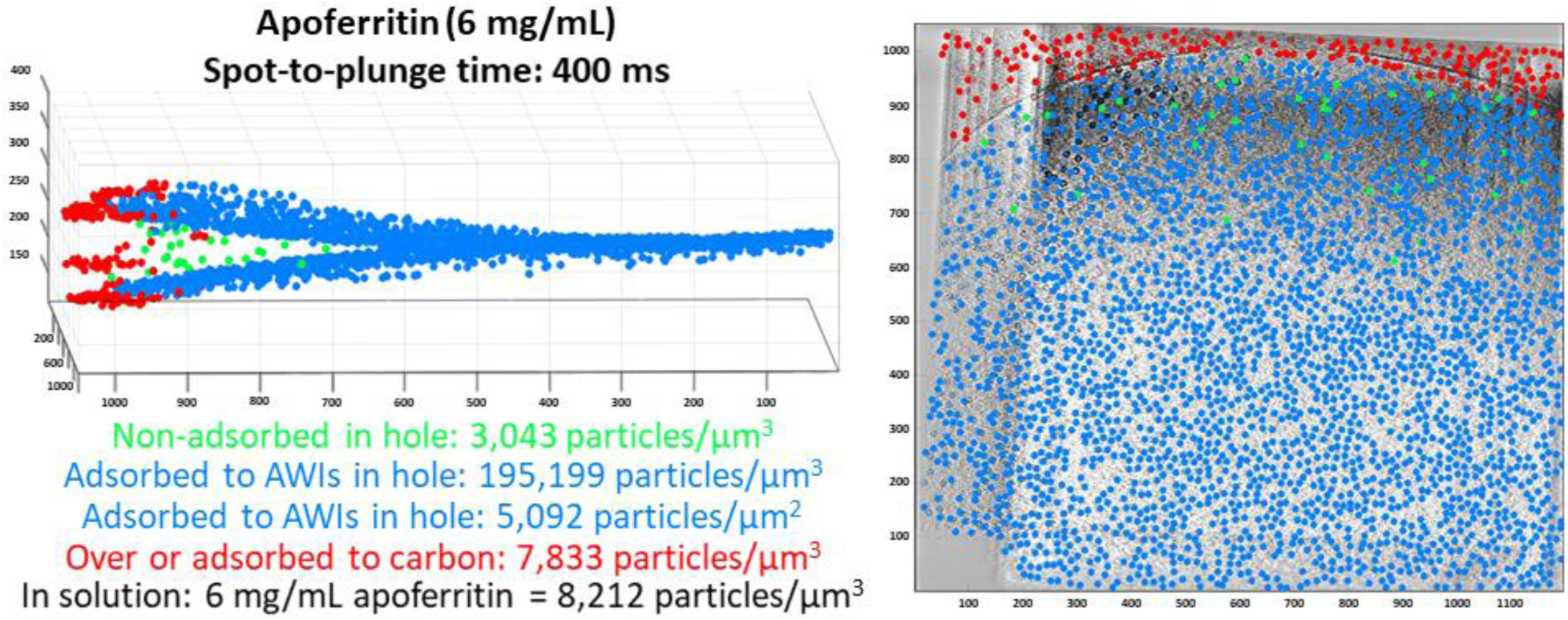
Particle distribution and concentrations of apoferritin (6 mg/mL = 8,212 particles/μm^3^ in solution) prepared using Spotiton and nanowire grids with spot-to-plunge time of 400 ms. Left depiction is a side-view aligned roughly perpendicular to the air-water interfaces. Right depiction is a top-down view with a tomographic slice included. Particles were manually picked according to their estimated locations relative to the air-water interface and carbon. Calculated particle densities (particles/μm^3^) are: 3,043 in the hole, 195,199 adsorbed to the air-water interfaces, and 7,833 over or adsorbed to carbon. Particle surface density at the air-water interface is 5,092 particles/μm^2^. Particle concentration calculations are estimated to be accurate to within 5%. Scale: 100 pixels is ~86 nm.

**Supplementary Figure 6.**
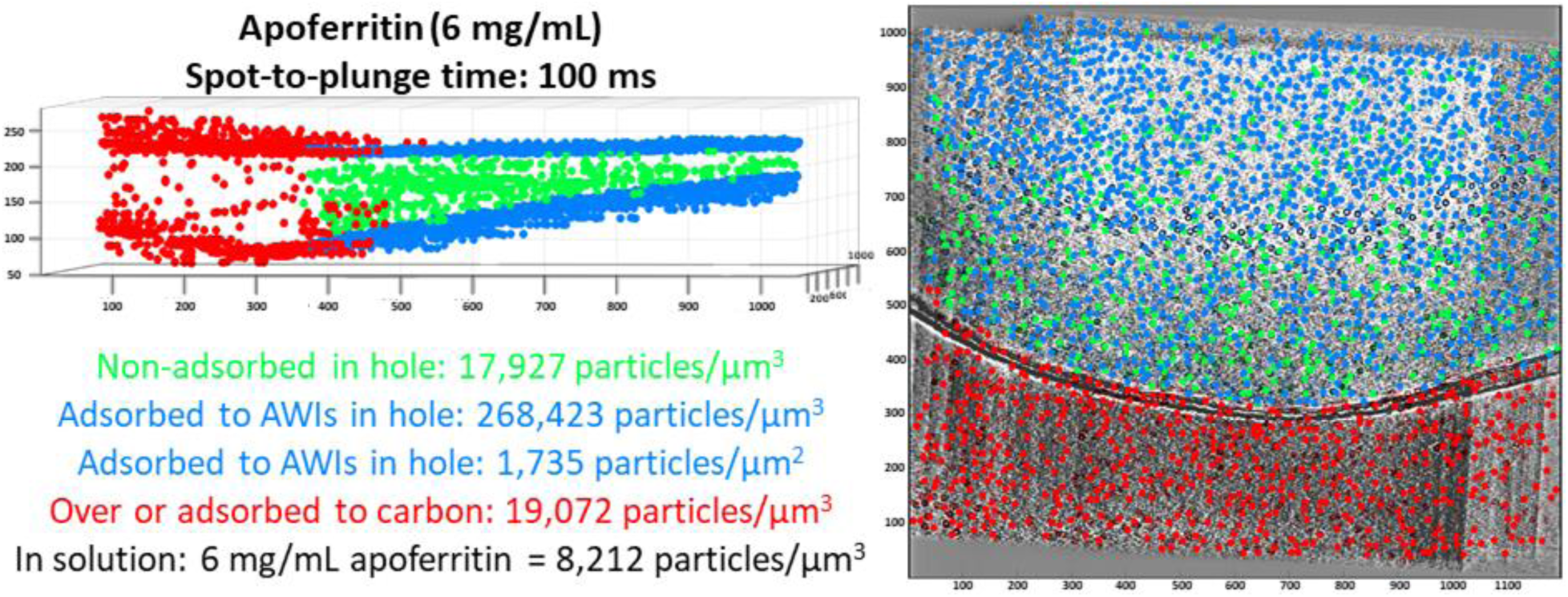
Particle distribution and concentrations of apoferritin (6 mg/mL = 8,212 particles/μm^3^ in solution) prepared using Spotiton and nanowire grids with spot-to-plunge time of 100 ms. Left depiction is a side-view aligned roughly perpendicular to the air-water interfaces. Right depiction is a top-down view with a tomographic slice included. Particles were manually picked according to their estimated locations relative to the air-water interface and carbon. Calculated particle densities (particles/μm^3^) are: 17,927 in the hole, 268,423 adsorbed to the air-water interfaces, and 19,072 over or adsorbed to carbon. Particle surface density at the air-water interface is 1,735 particles/μm^2^. Particle concentration calculations are estimated to be accurate to within 5%. Scale: 100 pixels is ~86 nm.

## Supplementary Videos

**Supplementary Video 1** | Tomogram slice-through of hemagglutinin prepared using Spotiton and nanowire grids with spot-to-plunge time of 800 ms, alongside a cross-sectional depiction. The vast majority of particles are observed to be adsorbed to the air-water interfaces with severe preferred orientation. Slice-through videos were rendered using ‘Slicer’ 3dmod from the IMOD package^36^ by orienting one plane of particles at the air-water interface to be roughly parallel to the field of view.

**Supplementary Video 2** | Tomogram slice-through of insulin-bound insulin receptor prepared using Spotiton and nanowire grids with spot-to-plunge time of 600 ms, alongside a cross-sectional depiction. The vast majority of particles are observed to be adsorbed to the air-water interfaces.

**Supplementary Video 3** | Tomogram slice-through of apoferritin with 0.5 mM TCEP (15 mg/mL) prepared using Spotiton and nanowire grids with spot-to-plunge time of 500 ms, alongside a cross-sectional depiction. Most of particles are observed to be adsorbed to the air-water interfaces, with a few non-adsorbed particles.

**Supplementary Video 4** | Tomogram slice-through of hemagglutinin prepared using Spotiton and nanowire grids with spot-to-plunge time of 100 ms, alongside a cross-sectional depiction. A majority of the particles are observed to be adsorbed to the air-water interfaces.

**Supplementary Video 5** | Tomogram slice-through of insulin-bound insulin receptor prepared using Spotiton and nanowire grids with spot-to-plunge time of 200 ms, alongside a cross-sectional depiction.

**Supplementary Video 6** | Tomogram slice-through of apoferritin with 0.5 mM TCEP (15 mg/mL) prepared using Spotiton and nanowire grids with spot-to-plunge time of 170 ms, alongside a cross-sectional depiction. Approximately nineteen times as many non-adsorbed particles are in the field of view as compared to the spot-to-plunge time of 500 ms (**Supplementary Video 3**).

**Supplementary Video 7** | Tomogram slice-through of apoferritin (6 mg/mL) prepared using Spotiton and nanowire grids with spot-to-plunge time of 400 ms, alongside a cross-sectional depiction. Most of the particles are observed to be adsorbed to the air-water interfaces, with a few non-adsorbed particles.

**Supplementary Video 8** | Tomogram slice-through of apoferritin (6 mg/mL) prepared using Spotiton and nanowire grids with spot-to-plunge time of 100 ms, alongside a cross-sectional depiction. Approximately six times as many non-adsorbed particles are in the field of view as compared to the spot-to-plunge time of 400 ms (**Supplementary Video 7**).

